# A TRAPPC2L/TRAPPC11/12/13 subcomplex directs TRAPPIII to autophagy

**DOI:** 10.64898/2025.12.02.691774

**Authors:** Mario Pinar, Vivian de los Ríos, Silvia Rodríguez-Pires, Eduardo A. Espeso, Miguel A Peñalva

**Author notes:** corresponding author at Department of Molecular Biosciences CSIC Centro de Investigaciones Biológicas Margarita Salas. Ramiro de Maeztu, 9 Madrid 28040, Spain.

## Abstract

TRAPP complexes are master regulators of membrane trafficking. TRAPPs are targeted to different locales by pathway-specific subunits decorating a core hetero-heptamer to build TRAPPII (Golgi exit) and TRAPPIII (autophagosomes and ER-Golgi traffic). Metazoan and Arabidopsis TRAPPIII have three components, TRAPPC11/12/13 absent from the budding yeast. We studied TRAPPC11/12/13 in the related ascomycete *Aspergillus nidulans*, where TRAPPC11 and TRAPPC12 localize to pre-autophagosomes and their ablation impairs autophagy. Two stable subcomplexes containing Tca17/TRAPPC2L coexist, one including the TRAPPII-specific subunits Trs120/Trs130/Trs65 and another including the TRAPPIII-specific subunits TRAPPC11/12/13. Both are recruited to core TRAPP by Tca17/TRAPPC2L, which therefore plays a crucial role by determining the fate of TRAPP. TRAPPIII also comes in two versions, TRAPPIIIa and TRAPPIIIb, both containing Trs85/TRAPPC8, but only TRAPPIIIb containing TRAPPC11/12/13, which target TRAPPIII to autophagy. This study might help characterize potentially pathogenic mutations affecting human TRAPPC11/12/13, facilitating assessment of their functional consequences in a genetically amenable ascomycete.

## Introduction

Rab GTPases regulate organelle identity by recruiting to membranes cytosolic proteins that determine the characteristic composition of each subcellular compartment ^1^. RABs are molecular timers alternating between active, GTP-loaded, and inactive, GDP-loaded, conformations, of which only the former exposes and inserts the prenylated C-terminus into a membrane and binds its cognate effectors efficiently ^2,3^. The high affinity that GDP has for the active site implies that the exchange of GTP for GDP requires the action of guanine nucleotide exchange factors (GEFs), which facilitate the release of GDP, making the nucleotide pocket accessible to GTP ^3,4^. It is broadly accepted that the membrane specificity of RABs is determined by the cognate GEF. Therefore, understanding the molecular physiology of these GEFs is a fundamental issue in cell biology.

TRAPPs (TRAnsport Protein Particle complexes) mediate nucleotide exchange on the GTPases RAB1 and RAB11 ^5–14^. Several TRAPP complexes have been described in the literature, all consisting of a core hetero-heptamer that is targeted to RAB1 a RAB11 depending on the presence of additional accessory TRAPP components. Intensive research has shaped the currently accepted paradigm according to which there are only two TRAPP complexes, denoted TRAPPII and TRAPPIII, TRAPPII is targeted to RAB11 by 3-4 subunits that modify the core, whereas TRAPPIII is targeted to RAB1 by Trs85/TRAPPC8 ^8–10,14–21^ (the composition of TRAPPs and the nomenclature of the different subunits is described in figure 1A).

**Figure 1:**
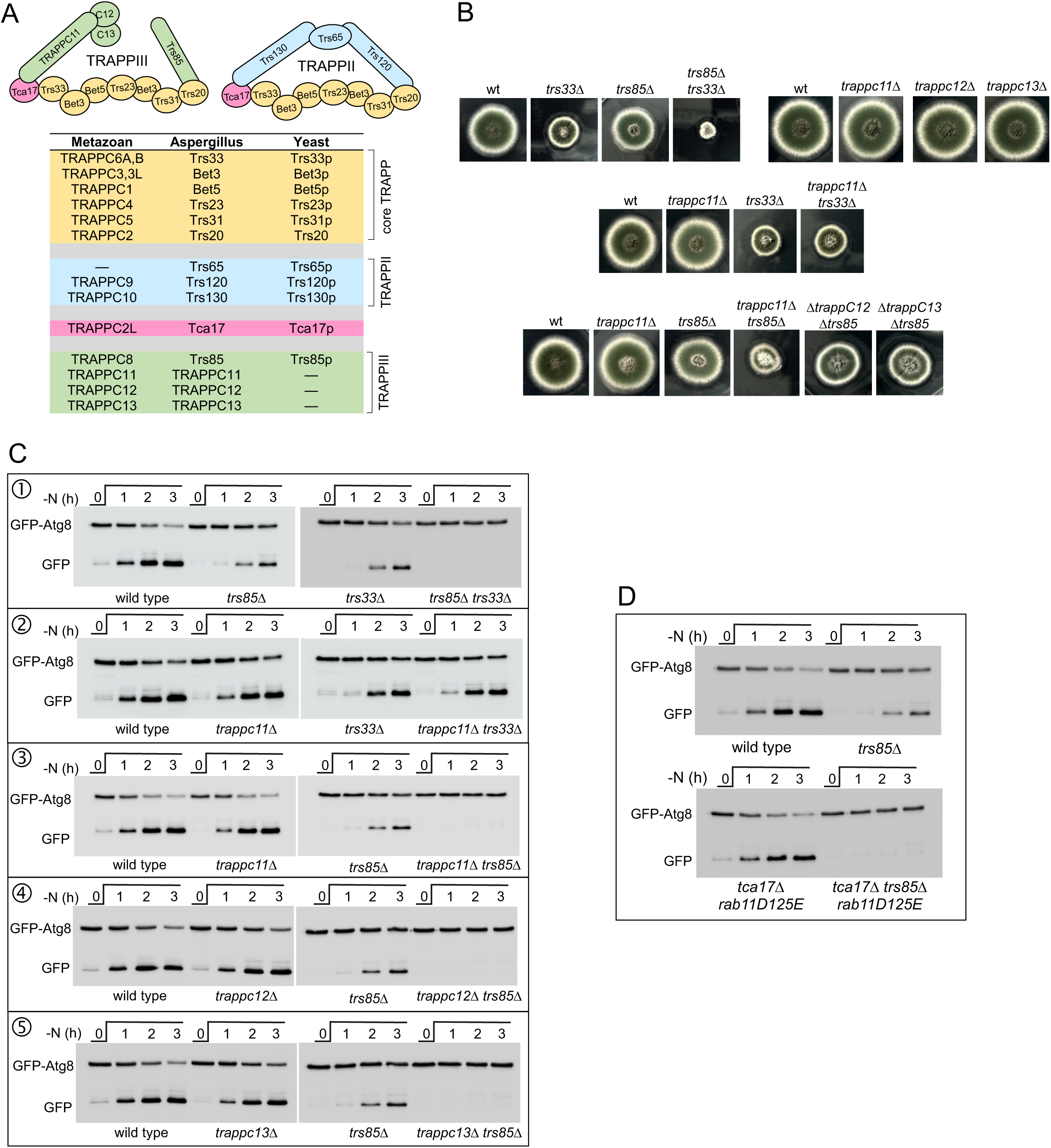
Ablation of Trs85 is insufficient to block autophagy. (A) Nomenclature of the proteins composing the fungal and metazoan TRAPP complexes and schematic representation of *A. nidulans* TRAPPII and TRAPPIII. (B) Growth tests of the indicated *A. nidulans* strains cultured on complete medium for three days at 37°C. (C) GFP-Atg8 processing assays in cells of the indicated genotypes that had been shifted from nitrogen-sufficient medium to nitrogen-starved medium to induce autophagy. In these assays, GFP-Atg8 present in the membrane of autophagosomes is delivered to the lumen of the vacuoles and proteolytically processed until the GFP moiety accumulates due to its resistance to vacuolar proteases. Full-length GFP-Atg8 and ‘free’ GFP were detected by anti-GFP Western blotting. (D) GFP-Atg8 processing assays demonstrating that autophagy is abolished in a *tca17*Δ *trs85*Δ double mutant.

RAB1, the substrate of TRAPPIII, participates in two physiologically antithetic pathways, exocytosis, specifically during transport between the ER and early Golgi, where RAB1 is recruited to COPII vesicles, and autophagy, where RAB1 is recruited to pre-autophagosomes ^10,13,17,18,22^. As yet outstanding is the problem of how TRAPPIII can differentially recruit and activate RAB1 to each of these locales. While a substantial amount of data on disease-causing amino acid substitutions in the human TRAPP components has accumulated ^23^, the only structure available to date for a metazoan TRAPPIII is that from *Drosophila melanogaster* ^11^. Studies with *Saccharomyces cerevisiae* have made key contributions to our understanding of the structural organization and physiology of the TRAPP complexes, but this unicellular eukaryote lacks homologues of TRAPPC11, TRAPPC12 and TRAPPC13, three of the eleven subunits of metazoan and plant TRAPPIII ^12,24,25^, thereby limiting further insight into mammalian TRAPPs exploiting this unicellular ascomycete. The genome of *S. cerevisiae* (12 Mbp and 5,900 coding genes) was shaped by reductive evolution resulting in extensive gene loss ^26^, a process that has not occurred in *Aspergillus nidulans* (30 Mbp and 12,000 genes), another genetically amenable ascomycete.

Using *A. nidulans* we determined how TRAPPII is directed towards RAB11 and characterized the composition of TRAPPs ^27,28^. We showed that unlike *S. cerevisiae*, *A. nidulans* has homologues of TRAPPC11/12/13 and that TRAPPIII is composed of the core hetero-heptamer plus Trs85 (TRAPPC8 in metazoans), like in the budding yeast ^27^. Here we revisit this conclusion by demonstrating the existence of two different TRAPPIII complexes, containing (TRAPPIIIb) or not (TRAPPIIIa, formerly TRAPPIII) TRAPPC11, TRAPPC12 and TRAPPC13, and establish that TRAPPC11, the subunit that links this subcomplex to the core, targets TRAPPIII to autophagy.

## Results

### Blocking autophagy requires ablation of both Trs85/TRAPPC8 and TRAPPC11/12/13

We reported previously that *A. nidulans* TRAPPIII, like its *S. cerevisiae* counterpart, consists of the banana-shaped core heterohexamer of essential subunits flanked at each end by a non-essential subunit, Trs33/TRAPPC6 and Trs85/TRAPPC8, respectively (Figure 1A) ^18,27^. Filamentous fungi grow by apical extension, a mechanism which is exquisitely dependent on exocytosis ^29^. Defects in exocytosis can be easily scored with growth tests monitoring colony size. A paradoxical observation was that *trs85*Δ impairs growth only slightly ^28^ (Figure 1B, top left), despite the fact that TRAPPIII is known to activate RAB1 during ER-to-Golgi transport ^10^. The second TRAPPIII non-essential subunit, Trs33 is also a component of TRAPPII (the GEF for RAB11 at the TGN), and therefore plays a dual exocytic role, which is reflected in the noticeable growth inhibition resulting from its ablation (Figure 1B, top left). We reasoned that sensitizing cells with a *trs33*Δ mutation could uncover the role of Trs85 in exocytosis. This was indeed the case. A double *trs33*Δ *trs85*Δ mutation showed a strong synthetic negative interaction (Figure 1B, top left).

In addition to its exocytic role, Trs85/TRAPPC8 is necessary and sufficient to target TRAPPIII to pre-autophagosomes (PASs), the Atg8-containing structures that give rise to autophagosomes, since *S. cerevisiae trs85*Δ precludes N-starvation-induced autophagy ^30^. However, mutant *Aspergillus trs85Δ* hyphae still retain significant Atg8 processing activity, a proxy of autophagy ^31^ (Figure 1C-1), establishing that there exist Trs85-independent mechanisms activating RAB1 in PASs. Notably, ablation of Trs33 impaired Atg8 processing to a similar extent than *trs85*Δ (Figure 1C-1), suggesting that in *Aspergillus* Trs33 cooperates with Trs85 in autophagy. Indeed, Atg8 processing was completely blocked in a *trs85Δ trs33Δ* double mutant strain (Figure 1C-1). Thus, *A. nidulans* Trs33 plays a direct or indirect role in autophagy.

Previously, we identified TRAPPC11/12/13 as minor components of TRAPPIII ^27^. However, mutant *trs33*Δ TRAPPIII does not contain TRAPPPC11/12/13 ^27^, indicating that these three subunits associate with TRAPPIII by way of Trs33 (or Tca17 or both) (Figure 1A). TRAPPC11/12/13 do not appear to have any exocytic role, as the corresponding single mutants grow like wild-type (Figure 1B, top right), and in double mutants *trs33*Δ and *trappc11*Δ do not show any synthetic negative interaction (Figure 1B, middle). Therefore, we hypothesized that these three subunits might be responsible for the autophagy deficit displayed by *trs33Δ* cells. We started by testing the effects of ablating TRAPPC11, because TRAPPC12 and TRAPPC13 are recruited to TRAPPIII by way of TRAPPC11 ^11^, and TRAPPC11, TRAPPC12 and TRAPPC13 form a stable subcomplex (see below). Indeed, GFP-Atg8 processing assays showed that ablating TRAPPC11 alone or in combination with *trs33*Δ impairs N-starvation-induced autophagy to a similar extent than *trs33*Δ (Figure 1C-2). In sharp contrast ablation of TRAPPC11 completely prevented it when combined with ablation of Trs85, mimicking the phenotype of *trs33*Δ *trs85*Δ double mutants (Figure 1C-3). Similar results were obtained after ablating TRAPPC12 and TRAPPC13, alone or in combination with *trs85*Δ (Figure 1C-4 and 1C-5). Therefore, TRAPPC11/12/13 play a physiological role in autophagy that accounts for the role of Trs33, explaining why *trs85*Δ alone does not fully prevent it. In conclusion efficient functioning of autophagy necessitates the concerted action of TRAPPC11/12/13 and Trs85. Notably, whereas single *trappc11*Δ, *trappc12*Δ, and *trappc13*Δ do not show any growth defect by themselves, *trappc11Δ*, but not *trappc12Δ* or *trappc13*Δ, reduces colony growth markedly when combined with *trs85*Δ (Figure 1B, bottom), suggesting that TRAPPC11 can take over, at least partially, the role of Trs85 in the ER-Golgi interface.

**Figure 2.**
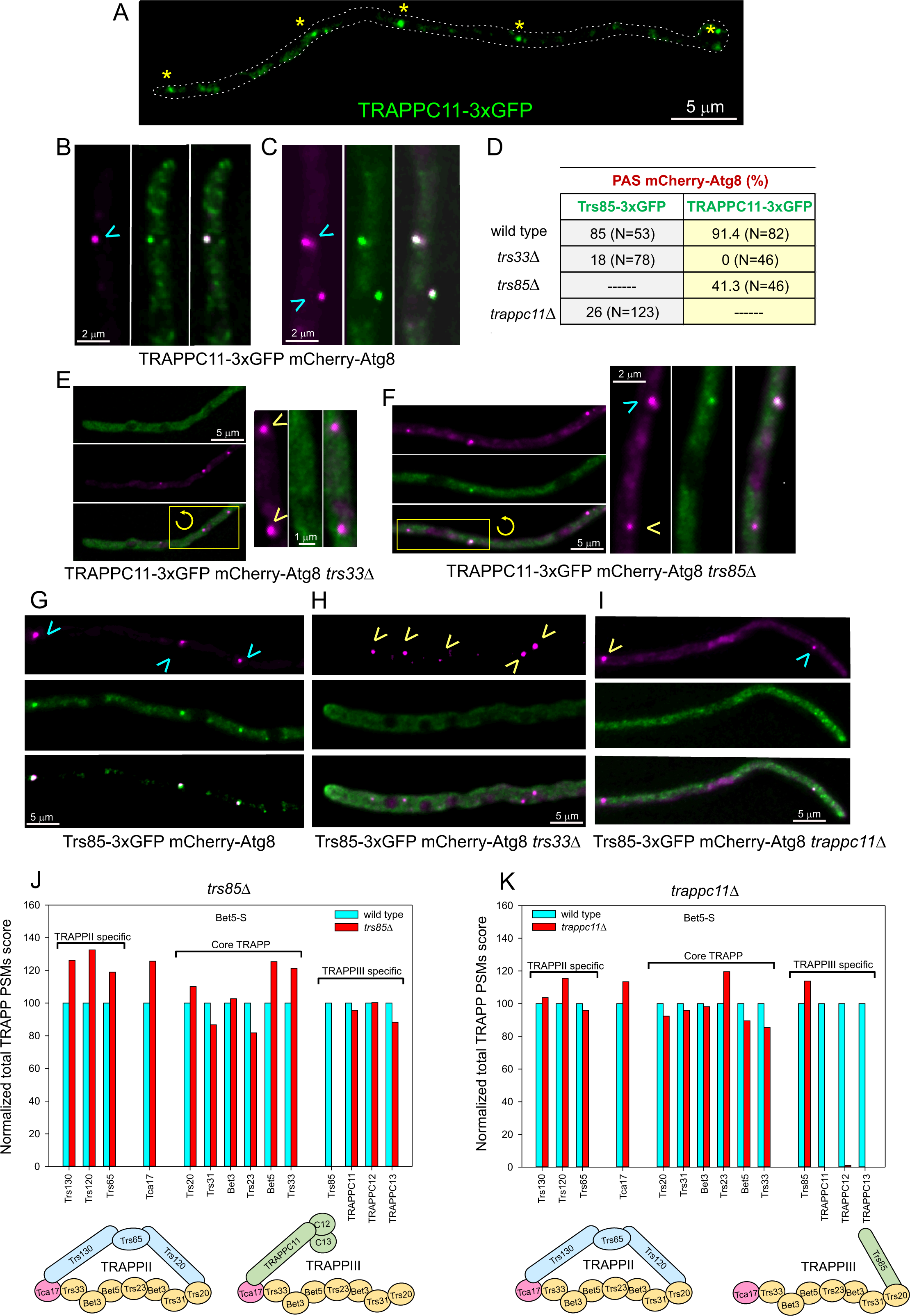
Endogenously tagged TRAPPC11 localizes to pre-autophagosomes. (A) localization of TRAPPC11 to pre-autophagosomes across the length of a hypha. (B, E-I) PASs are labeled with mCherry-Atg8. PASs containing TRAPPC11 are indicated with cyan arrowheads, whereas those that do not contain TRAPPC11 are indicated with yellow arrowheads. (B) TRAPPC11 colocalizes with a PAS near the tip. (C) TRAPPC11 is present in PASs located away from the tip. (D) Quantitative data for localization experiments. (E) TRAPPC11 does not localize to the PAS in cells ablated for Trs33. (F) The number of PASs containing TRAPPC11 is reduced in cells lacking Trs85. (G) Trs85 localizes to PASs. (H) Trs85 localization to PASs is largely impaired in cells lacking TRS33. (I) TRP85 localization to PASs is impaired in cells lacking TRAPPC11. (J) The composition of TRAPPs, purified from *trs85*Δ cells by Bet5-S affinity chromatography, was analyzed by MS/MS. The actual PSM counts for each protein in the wild-type condition was set as 100%. (K) As in (J) for *trappc11*Δ cells.

**Figure 3.**
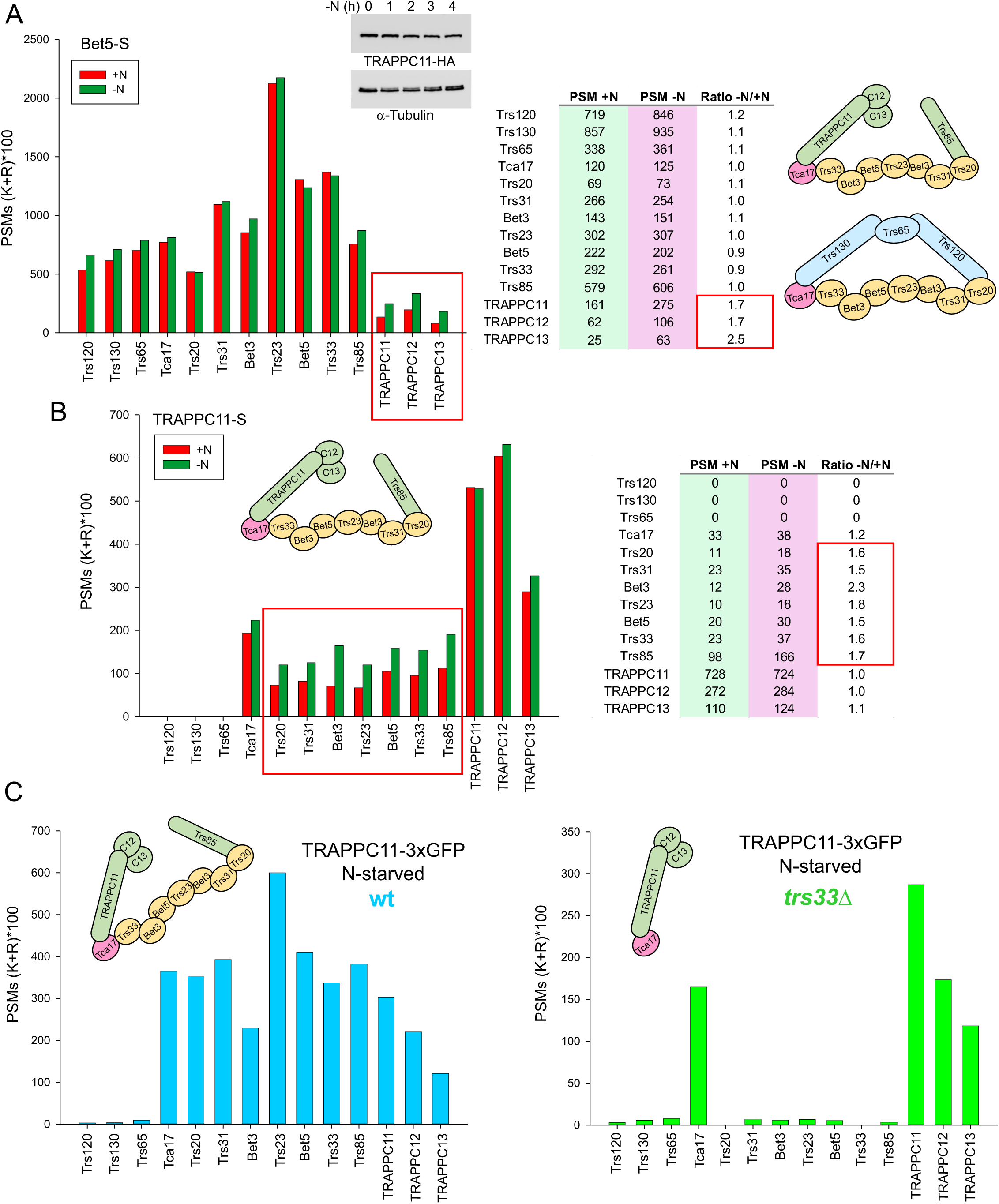
The amount of core TRAPP associated with a subcomplex containing TRAPPC11/12/13 and Tca17/TRAPPC2L increases upon shifting cells to N-starved conditions. (A) Abundance of TRAPP subunits in Bet5-S purified material. Bet5 is a pan-TRAPP subunit. Bars indicate spectral counts normalized to the number of Lys and Arg residues for each protein (which is used as an approximation to the number of tryptic peptides potentially detectable by MS/MS for each protein) multiplied by 100. Displayed on the right table are the ratios of actual spectral counts obtained under N-sufficient (N+) and N-starving (N-) conditions. The western blot in the middle shows that the amount of HA_3_-tagged TRAPPC11 does not increase after starving cells for nitrogen (anti-alpha-tubulin was used as a loading control). (B) the amount of TRAPP core subunits associated with TRAPPC11 increases upon shifting cells to nitrogen-starved conditions. Note that TRAPPC11 does not associate at all with TRAPPII-specific subunits Trs120, Trs130, and Trs65, but pulls-down Tca17, a subunit reportedly associated with TRAPPII. The table on the right indicates relative values obtained by comparing the ratio between N-sufficient (N+) and N-starving conditions (N-). (C) A stable subcomplex containing Tca17 and TRAPPC11/12/13 can be isolated from cells lacking Trs33. GFP-trapping experiments in which proteins were immunoprecipitated with anti-GFP magnetic agarose beads and subjected to MS/MS analysis.

**Figure 4.**
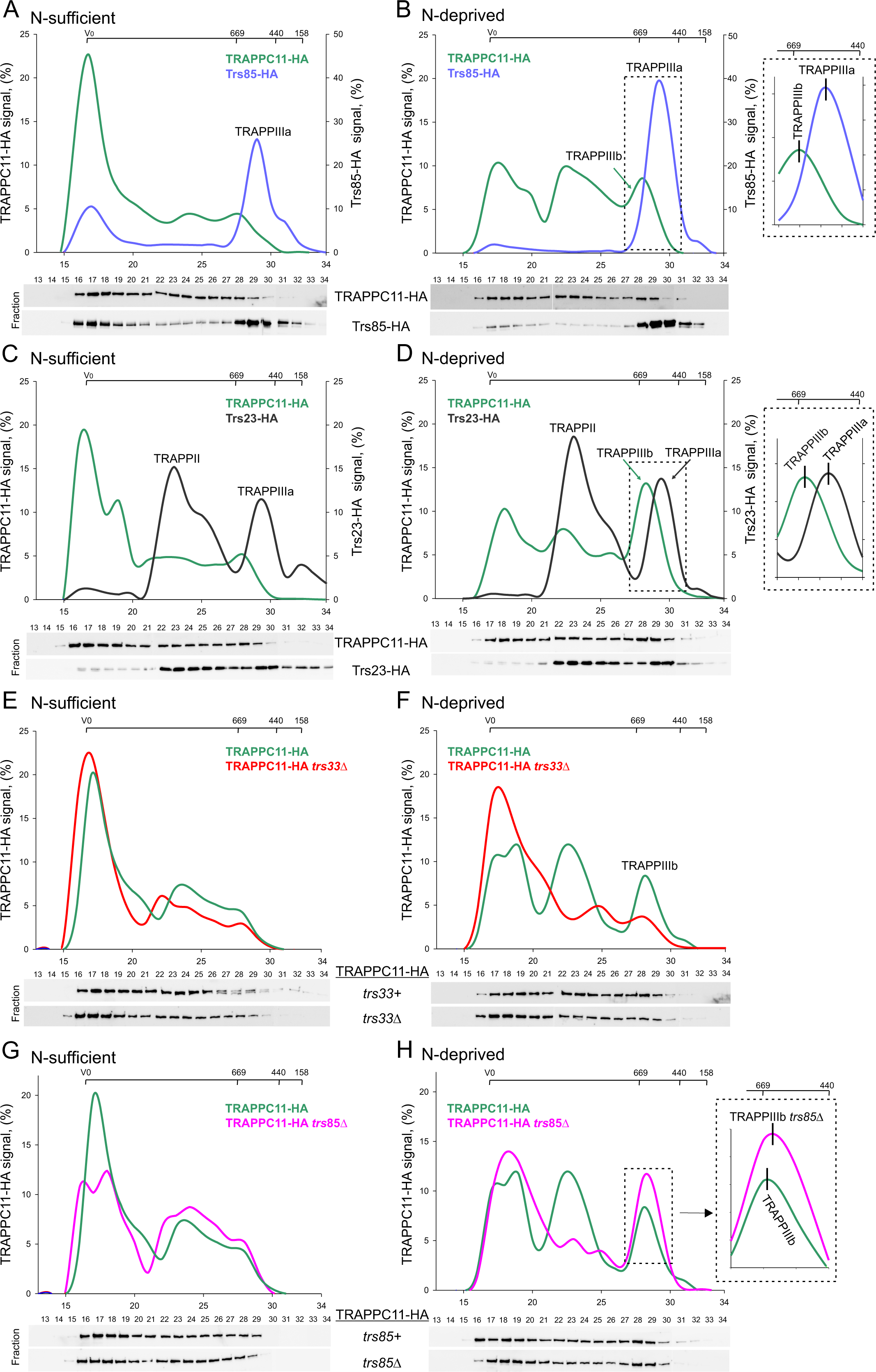
Two versions of TRAPPIII, containing or not TRAPPC11/12/13 are present in *A. nidulans* extracts. (A-B) Extracts of cells co-expressing HA_3_-tagged versions of TRAPPC11 and Trs85 cultured under nitrogen-sufficient (A) and nitrogen-deprived conditions (B) were run through a Superose 6 column and the resulting fractions analyzed by anti-HA western blotting. TRAPPC11, is with a complex that we denote TRAPPIIIb whose relative abundance increases upon nitrogen starvation and that eludes ahead of TRAPPIII labeled with Trs85, that we denote TRAPPIIIa. (C-D) As in (A-B) but using HA_3_-tagged core subunit Trs23 instead of Trs85 to mark the position of all TRAPP complexes. (E-F) the assembly of TRAPPIIIb is dependent on Trs33. (G-H) TRAPPIIIb does not dissemble in absence of Trs85, but its elution is delayed, demonstrating that it contains Trs85.

### Efficient recruitment of TRAPPIII to pre-autophagosomes requires both TRAPPC11 and Trs85

Consistent with its proposed role in autophagy, subcellular localization studies of TRAPPC11 endogenously tagged with triplicated GFP showed that it localizes to puncta, a localization typical of *A. nidulans* pre-autophagosomes (PASs) (Figure 2A) ^31^. Indeed, these puncta colocalized with the prototypic PAS marker Atg8 (Figure 2B-C). Like TRAPPC11, TRAPPC12 also localizes to PAS structures (Figure S1) (TRAPPC13 was undetectable after endogenous tagging). As expected from the recruitment of TRAPPC11(/12/13) to TRAPPIII being mediated by Trs33, TRAPPC11-3xGFP localization to the PAS is dependent on the presence of the latter (Figure 2D). Thus, 91% of TRAPPC11 puncta contain Atg8 in the wild-type, contrasting with none in *trs33*Δ (Figure 2D-E).

Notably, subcellular localization experiments supported the view that Trs85 and TRAPPC11 cooperate to target TRAPPIII to the PAS: Less than half (41%) of Atg8 PASs contain TRAPPC11 in *trs85*Δ hyphae (Figure 2D and F), only 18% (N=78) of the Atg8 PASs displayed detectable levels of Trs85-3xGFP in *trs33*Δ cells compared to 85% (N=53) in the wild-type (Figure 2D, G-H); and only 26% of Atg8 PASs contained Trs85-3xGFP in *trappc11Δ* cells (Figure 2D and I). As neither *trs85*Δ or *trappc11Δ* disorganizes TRAPPIII (Figure 2J and 2K) (ref. ^27^ for *trs33*Δ), it follows that the efficient recruitment of TRAPPIII to PASs necessitates both Trs85 and TRAPPC11, explaining the partial defect in Atg8 processing documented in the absence of either.

### TRAPPC11/12/13 associate with Tca17/TRAPPC2L in a stable subcomplex

MS/MS analysis of the composition of TRAPPIII pulled down by S-tagged Trs85 had revealed that TRAPPC11/12/13 subunits are sub-stoichiometric ^27^, but this analysis had been carried out with cells cultured under N-sufficient conditions. Therefore, we compared the composition of TRAPP complexes isolated from cells cultured under N-sufficient and N-starving conditions. To this end, we purified TRAPPs by S-agarose affinity with S-tagged Bet5/TRAPPC1, a pan-TRAPP subunit ^27^. Notably, the amount of TRAPPC11/12/13 associated with TRAPP augmented markedly when cells were shifted to nitrogen-starving conditions (Figure 3A), which occurred despite the fact that levels of TRAPPC11 did not increase (Figure 3A, middle top). As none of the other TRAPP components increased its association with the complex, these data strongly indicate that N-starvation stimulates the incorporation of TRAPPC11/12/13 onto TRAPPIII, contributing to target TRAPPIII to autophagy. MS/MS data in Figure 3A also indicate that TRAPPC11/12/13 are sub-stoichiometric compared to Trs85, even under nitrogen-starving conditions, suggesting that a proportion of TRAPPIII does not contain them.

To buttress the conclusion that N-starvation stimulates the incorporation of TRAPPC11/12/13 onto TRAPPIII, we purified TRAPPs from cells expressing TRAPPC11-S-tag. These experiments reinforced the conclusion that the amount of core TRAPP associated with TRAPPC11 increases when cells were cultured under nitrogen-starving conditions (Figure 3B). MS/MS analysis of the composition of TRAPP complexes additionally provided a key piece of information, namely that TCA17/TRAPPC2L, a component of *Aspergillus* TRAPPII, copurifies with TRAPPC11, which is in agreement with the *Drosophila* structure establishing that TRAPPC11 is recruited to TRAPPIII by way of TRAPPC2L ^11^. Therefore, our data suggest the existence of two TRAPPIII complexes, one containing core TRAPP plus Trs85, denoted TRAPPIIIa and a second containing, in addition, Tca17 and TRAPPC11/12/13, denoted TRAPPIIIb. The finding that a fungal TCA17 is a component of TRAPPIII is notable because TCA17 is absent from *S. cerevisiae* TRAPPIII ^13,17,18,32^.

In previous work, we identified a stable TRAPPII subcomplex consisting of Trs120, Trs130, Trs65 and Tca17 that can be assembled onto the core TRAPP hetero-heptamer to generate TRAPPII ^27^. To test whether TRAPPC11/12/13 act in a complex we purified the proteins associated with TRAPPC11-3xGFP in cells lacking Trs33, which disengages them from core TRAPP. This experiment (Figure 3C) revealed that the only proteins copurifying with TRAPPC11 were TRAPPC12, TRAPPC13 and Tca17, indicating that they form a stable subcomplex. This is a remarkable finding, as it implicates that Tca17 (TRAPPC2L) determines the fate of the TRAPP core by recruiting the TRAPPII-specific subcomplex or the TRAPPIIIb specific subcomplex to core TRAPP.

As *Aspergillus* Trs33 recruits TRAPPC11/12/13 by way of Tca17, ablation of Tca17 should phenocopy the consequences of removing TRAPPC11/12/13 on autophagy by impeding the assembly of the subcomplex that incorporates them into TRAPPIII. Tca17 is essential for viability because its ablation disorganizes TRAPPII, preventing exocytosis^27^. However, the role of Tca17 in autophagy can be assessed after bypassing the role of TRAPPII with a D125E missense mutation that constitutively activates RAB11, thereby rescuing viability ^8,27^. *rab11^D125E^ tca17*Δ mutants display negligible impairment of autophagy, whereas *rab11^D125E^ tca17*Δ *trs85*Δ triple mutants show a complete blockage (Figure 1D). These data strongly support a dual role of Tca17 in both autophagy and exocytosis by way of its ability to scaffold TRAPPII- and TRAPPIIIb-specific subcomplexes.

Two regions of *D. melanogaster* TRAPPC11 (residues 442 through 453 and 478 through 486) within a 15-alpha-helix solenoid that comprises most of N-terminal half of the protein interact with an alpha-helix in TRAPPC2L that contains Leu-Asp-Φ motif ^11^ (Φ indicates a hydrophobic residue). These features are conserved in both humans, *Arabidopsis* and *Aspergillus* (Figures S2-S3). Indeed, AlphaFold predicts that the TRAPPC11 15-helix solenoid and its interaction with the Leu-Asp-Φ motif in Tca17 (TRAPPC2L) is conserved (Figure S3). Density maps of the available TRAPPIII structure did not allow modelling of the C-terminal half of TRAPPC11, nor of TRAPPC12/TRAPPC13 ^11^. Thus, we run AlphaFold predictions with these polypeptides, using the *Aspergillus*, *Drosophila* and human sequences. These three predictions yielded remarkably similar models (Figure S4A). In all three cases the C-terminal half of TRAPPC11 consisted of three consecutive β-sandwiches, of which the most C-terminal interacted with both TRAPPC12 and TRAPPC13. TRAPPC13 consisted of two β-sandwiches in *Aspergillus* and three in humans and *Drosophila* (Figure S4B). TRAPPC12 is a sixteen-helix α-barrel. The two beta sandwiches of *Aspergillus* TRAPPC13 and its equivalents in the human and *Drosophila* predicted protein structures are docked against TRAPPC12. The β-sandwiches are separated by an ∼ 50 residue loop that passes through the central cavity of the TRAPPC12 α-barrel, contributing to the intimate association between these two subunits (Figure S4B-C).

### Analysis of TRAPPs under N-sufficient and N-starved conditions support the existence of two TRAPPIII complexes

To investigate the recruitment of TRAPPC11/12/13 to TRAPPIII, we compared the elution profiles of TRAPPC11-HA_3_ and Trs85-HA_3_ after chromatographing cell-free extracts of hyphae co-expressing the respective tagged proteins through a gel filtration column and determining by anti-HA Western blotting the amount of Trs85 and TRAPPC11 in each fraction (Figure 4A). The predicted *Mr* of TRAPPIIIa (core TRAPP plus Trs85) is 237 kDa, whereas that of TRAPPIIIb (TRAPPIIIa plus TRAPPC11/12/13 and Tca17) is 485 kDa. Because TRAPPIIIa is rod-shaped, it elutes between the 669 kDa and 440 kDa standards ^27^.

Under nitrogen-sufficient conditions most of TRAPPC11 eluted with the void volume, whereas Trs85 was resolved as a single peak corresponding to TRAPPIIIa (Figure 4A). In contrast, with N-starved cells, a peak of TRAPPC11 was detected at the region of Trs85, although eluting ahead of it (Figure 4B). We interpreted that this peak containing both TRAPPC11 and Trs85 (Figure 4B blots) corresponds to TRAPPIIIb. To buttress this interpretation, we chromatographed extracts expressing the core subunit Trs23, which marks the positions of all TRAPPs ^27^ (Figure 4C). In N-starved cells a large proportion of TRAPPC11 was shifted to this peak eluting ahead of TRAPPIIIa, whose position was now labelled with Trs23 (Figure 4D).

Two additional pieces of data contributed to the evidence that this forward-shifted TRAPPIII peak corresponds to TRAPPIIIb. One was that it disappeared when Trs33 was ablated, irrespective of the nitrogen condition (Figure 4E-F). The second was that it is present in cells ablated for Trs85, but its elution is slightly retarded as would be expected from the loss of Trs85 (Figure 4G-H, see box). In summary, these experiments established that inducing autophagy increased the abundance of TRAPPIII containing TRAPPC11/12/13 (i.e., TRAPPIIIb), in agreement with the above MS-MS analysis. Importantly, they strongly support the existance of two versions of TRAPPIII, including (TRAPPIIIb) or not (TRAPPIIIa) TRAPPC11/12/13. It is tempting to speculate that TRAPPIIIa plays a general role serving both the autophagy and secretory routes whereas TRAPPIIIb is specialized in autophagy.

## Discussion

The mechanistic bases of the autophagy pathway have been largely dissected through studies with *S. cerevisiae*. Subsequent work demonstrated that this pathway is conserved in metazoans, with a few exceptions. A notable example is that of the oligomeric complex TRAPPIII, which serves as a GEF for the RAB GTPase, RAB1/Ypt1, a master regulator of autophagy and exocytosis. How the same complex discriminates between two such opposing physiological roles (one anabolic, the second catabolic) has started to be understood only recently ^33^. One important caveat is that *S. cerevisiae* TRAPPIII differs from that in metazoans and plants, which contains three subunits denoted TRAPPC11, TRAPPC12 and TRAPPC13 that are absent in the former ^24,25,34^. This is relevant because mammalian TRAPPC11 has been shown to play a role in autophagy, mediated by its interaction with ATG2B, a lipid transfer protein which facilitates phagophore expansion ^35^. We have previously shown that the filamentous ascomycete *A. nidulans*, a close relative of *S. cerevisiae* that has not undergone a genomic reduction, is a useful model to study autophagy. For example, in higher cells autophagosomes are formed in the proximity of ER domains denoted omegasomes, which are absent in the budding yeast, but present in *Aspergillus* ^31^. We have also reported that RAB1, besides being present in secretory membranes, localizes to the PAS and the expanding phagophores ^31^ and that, in agreement, Trs85, a diagnostic subunit of TRAPPIII, is present both on Golgi membranes and in the PASs ^28^. Importantly, *A. nidulans* contains homologs of TRAPPC11, TRAPPC12, and TRAPPC13 ^27^.

We show that Tca17 and TRAPPC11/12/13 are components of a TRAPPIII complex targeted to the autophagy pathway. Efficient autophagy requires both these four components and Trs85. TRAPPC11/12/13 form a stable subcomplex with Tca17 (TRAPPC2L). Tca17 binds to Trs33 and Bet3a ^11,21,36^. This dual interaction of Tca17 with both Trs33 and Bet3a/TRAPPC3a explains that while *Aspergillus* Tca17/TRAPPC2L is essential for growth, Trs33/TRAPPC6 is not. Trs20 located at the TRAPPIII end opposite to Tca17 connects the core to Trs85/TRAPPC8, demonstrating that the organization of TRAPPIII in *A. nidulans* is akin to that in *D. melanogaster*, the only organism for which a three-dimensional structure is available ^11^. Importantly, our studies strongly suggest that there are two co-existing versions of TRAPPIII, both containing Trs85 but only TRAPPIIIb containing Tca17 and TRAPPC11/12/13, which are required to target efficiently the complex to the PAS. TRAPPIIIa, which does not contain these four proteins, would play its role in the secretory pathway.

Somewhat paradoxically, considering that fungal growth is exquisitely dependent on the efficient functioning of the secretory pathway, ablation of Trs85 results in a weak growth phenotype, despite the fact that TRAPPIII is involved in transport between the ER and the Golgi. Our finding that *trappc11*Δ shows a synthetic negative interaction with *trs85*Δ strongly suggests that in the absence of the latter, TRAPPC11, but not TRAPPC12 or TRAPPC13, are able to perform partially the role of Trs85. Trs85 anchors the complex to membranes ^13^. Thus, this interaction is consistent with the possibility raised by Galindo and Munro that TRAPPC11 shares with Trs85 the ability to anchor the complex to membranes ^24^.

Mutations affecting human TRAPPC11, TRAPPC12 and TRAPPC2L cause a complex spectrum of severe developmental and neurological disorders ^23,37–39^. An α-helix in fungal, plant and metazoan TRAPPC2L/Tca17 containing a conserved Leu-Asp-Φ motif it is crucial for its interaction with TRAPPC11. In humans, a homozygous missense mutation resulting in Asp37Tyr substitution within this motif causes a developmental disorder, but this mutation is relatively uninformative in terms of understanding the basis of the disease at the molecular level, as it affects both TRAPPII and TRAPPIII through interaction with TRAPPC10/Trs130 and TRAPPC11, respectively. In contrast, *A. nidulans* can be a useful model for the functional analysis of non-synonymous mutations in TRAPPC11, which might help to improve the prognosis of the spectrum of diseases caused by mutations in this gene.

The finding that two stable subcomplexes containing Tca17 are present in *Aspergillus* is important. One contains Trs120, Trs130, and Trs65 and when combined with core TRAPP gives rise to TRAPPII, targeting TRAPP to the TGN. The second contains Tca17, TRAPPC11, TRAPPC12, and TRAPPC13, and when combined with TRAPPIIIa gives rise to TRAPPIIIb, thereby targeting TRAPP to autophagosomes.

## Acknowledgements

This work has been supported by Grants TED2021-129607B-I00 funded by MCIN/AEI/ 10.13039/501100011033 and by the “European Union NextGeneration EU/PRTR”, by Grant PID2021-124278OB-I00 funded by MCIN/AEI/ 10.13039/501100011033/FEDER,UE and “ERDF A way of making Europe”, and by Grant PID2024-1-155595NB-I00 funded by MCIN/AEI/ 10.13039/501100011033/FEDER,UE, to EAE and MAP.

S.R-P is Staff hired under the Generation D initiative, promoted by Red.es, an organisation attached to the Ministry for Digital Transformation and the Civil Service, for the attraction and retention of talent through grants and training contracts, financed by the Recovery, Transformation and Resilience Plan through the European Union’s Next Generation funds.

## Supplemental Figure legends

**Figure S1.** Localization of GFP-tagged TRAPPC11 and TRAPPC12 to secretory compartments and to the PASs. The latter are indicated by arrowheads.

**Figure S2.** Four-way amino acid sequence alignment of human, *Drosophila*, *Arabidopsis* and *Aspergillus* TRAPPC11. The *foie gras* domain is shaded in purple. The two regions that Galindo et al determined as being involved in the interaction with TRAPPC2L are boxed in yellow. Similar amino acids were according to the Blosum 45 score table.

**Figure S3:** Top, four-way amino acid sequence alignment of *Aspergillus*, human *Drosophila* and *Arabidopsis* TRAPPC2L/Tca17. The region containing the critical Leu-Asp-Φ motif is boxed (Φ indicates a hydrophobic residue). Similar amino acids were according to the Blosum 45 score table. Bottom, AlphaFold prediction of the interaction between the *foie gras* domain of *Aspergillus* TRAPPC11 and Tca17. Left, overall view of the predicted complex. Right, details of predicted interactions between the relevant region of the TRAPPC11 *foie gras* domain with α-helix 1 in Tca17. Asp48 within the conserved Leu-Asp-Φ motif corresponds to Asp37 in both *Drosophila* and humans.

**Figure S4.** AlphaFold predictions of the complex formed by the C-terminal half of TRAPPC11 with TRAPPC12 and TRAPPC13. (A) Predictions for the indicated complexes shown in surface and cartoon representations. (B) Structural alignment of TRAPPC13 from the three species showing the loop that is located between two beta-sandwiches. Note that *Aspergillus* TRAPPC13 contains only two beta-sandwiches, whereas the human and *Drosophila* proteins contain three. (C) AlphaFold prediction of the loop of *Aspergillus* TRAPPC13 inserted through the central channel of the TRAPPC12 alpha barrel.

## Star Methods

### Experimental model and subject details

#### Aspergillus techniques

For strain propagation and harvesting of conidiospores we used standard *Aspergillus nidulans* media. Deletion and epitope-tagged alleles were introduced into appropriate genetic backgrounds by meiotic recombination and/or transformation. For latter, we used *nkuA*Δ strains deficient in the non-homologous end-joining pathway as recipients ^40^.

#### Null mutant strains and protein tagging

*trs33*Δ, *trs85*Δ, *trappc11*Δ, *trappc12*Δ, *trappc13*Δ were generated by gene replacement, using as donor DNA molecules cassettes constructed by fusion PCR ^41^. Integration events were confirmed by PCR with external primers. The following proteins were C-terminally tagged endogenously: Trs85-3xGFP ^28^, TRAPPC11-3xGFP, TRAPPC11-HA_3_, TRAPPC11-Stag, Bet5-stag, Trs85-HA_3_, and Trs23-HA_3_ ^8,27^. The oligonucleotides used to construct these cassettes are listed in Table S2. GFP-Atg8 and mCherry-atg8 were expressed under the control of *gpdA^mini^* promoter ^31^.

### Method details

#### Autophagy assays

For GFP-Atg8 processing experiments, carried out essentially as described ^31^, *A. nidulans* strains were cultured for 13–14 hours at 30°C on synthetic complete minimal medium supplemented with 1% (w/v) glucose as carbon source and 40 mM ammonium sulphate as nitrogen source. Mycelia were harvested by filtration and an adequate proportion was were freeze-dried (zero time point). The remaining mycelia were transferred to flasks containing the above medium without any nitrogen source, to induce autophagy. Mycelial samples were collected every hour for up to 3 hours and freeze-dried. 5 mg of each freeze-dried mycelial lyophilizate was ground to a fine powder with a ceramic ball using a FAStprep-24 (MP Biomedicals) and 2-ml screw-cap tubes. Total protein extracts were obtained by alkaline lysis ^42^. 10 µl of each sample were run through 12% SDS-polyacrylamide gels before being electro-transferred to nitrocellulose filters, which were reacted with α-GFP antibody (1:4000) and HRP-coupled anti-mouse (1:5000) as primary and secondary antibodies respectively. The protein signal was detected with Clarity western ECL substrate (Biorad) and the chemiluminescence was recorded with a Biorad Chemidoc Imaging System.

#### Determining levels of TRAPPC11 upon inducing autophagy

Conidiospores from a strain expressing TRAPPC11-HA_3_ were cultured for 14 hours at 30°C on synthetic complete minimal medium with 1% (w/v) glucose as carbon source and 20 mM ammonium sulphate as nitrogen source. Mycelia were harvested by filtration and transferred to the above medium without ammonium sulfate. Samples of mycelia were collected every hour for 4 hours; total proteins were extracted as described above and analyzed by western blotting. TRAPPC11-HA_3_ signal was detected with rat α-HA primary antibody (1:1000) and HRP-coupled-α-rat IgM plus IgG secondary antibody (1:4000). α-tubulin, used as a loading control, was detected with α-tubulin antibody (1:5000) and HRP-coupled-α-mouse antibody (1:4000).

#### Fluorescence microscopy

In vivo fluorescence microscopy experiments were carried out on µ-Slide 8 well uncoated chambers (Ibidi), with 0.3 mL of “watch minimal medium” (WMM) ^43^ per well, which were inoculated with conidiospores suspensions and incubated overnight at 26-28°C. The microscope setup consisted of a Leica DMI6000B inverted microscope, a Leica 63×/1.4 N.A. plan apochromatic objective coupled to an objective heater (Peacon) adjusted to 28°C and a Hamamatsu ORCA-ER CCD (1344×1024 pixels, 6.45 µm cell size) camera. The hardware was driven by Metamorph Premier software (Molecular Dynamics) and coupled Single-channel acquisition of fluorescence images was made with Semrock GFP-3035B and TXRED-4040B ‘BrightLine’ filter cubes. All colocalization experiments were made with images acquired simultaneously in the two channels with a Photometrics Dual-View beam splitter (CCD camera). Z-stacks of images acquired with a Z-pass of 0.5 µm were transferred to the computer RAM using the ‘stream acquisition’ function of Metamorph to minimize the time elapsed between acquisition of the first and last planes of the stack. Z-stacks were deconvolved using Huygens Professional software (Hilversum). Maximum Intensity Projections (MIPs) of the red and green channels were merged using the ‘color align’ Metamorph plug in. Images were annotated with Corel Draw.

#### Superose 6 size exclusion chromatography

Strains were cultured on *Aspergillus* fermentation medium (MFA) consisting of minimal medium supplemented with 2.5% (v/v) of corn steep liquor syrup (Solulys 048R, Roquette Laisa S.A., Valencia, Spain), 50mM each of Na_2_HPO_4_ and NaH_2_PO_4_, 50mM NaCl, 3% (w/v) sucrose as main carbon source, 20mM (NH4)_2_SO_4_ as main nitrogen source and vitamins as required to supplement the auxotrophic mutations present in each strain. After 13-14 hours of incubation at 30°C with shaking a mycelial sample was collected by filtration and freeze-dried (N-sufficient conditions). The rest of the mycelium was transferred to synthetic complete minimal medium without nitrogen and further incubated for 1 hour at 30°C before harvesting and freeze-drying the resulting biomass (N-starved conditions). 80 mg of each freeze-dried mycelial sample were ground in 2 ml crew-cap microcentrifuge tubes containing a ceramic bead with a 20 sec pulse of FastPrep set at power 4. The powder was resuspended in 1.5 ml of lysis buffer containing 25mM Tris-HCl, pH 7.5, 0.5% v/v NP40, 600mM KCl, 4mM EDTA, 1mM DTT, 5 μM MG132 and Complete ULTRA EDTA-free protease inhibitor cocktail (Roche). ∼ 100 μl of 0.6 mm glass beads were added to the suspension, which was homogenized with a 15 sec pulse at full power of the FastPrep followed by incubation for 10 min at 4°C in a rotating wheel. This step was repeated two additional times before the resulting homogenate was centrifuged at 15,000 x g and 4°C in a refrigerated microcentrifuge. 200 μl of the supernatant were loaded onto a Superose 6 10/300 column (Cytiva) equilibrated in lysis buffer without protease inhibitors, which was run at a flow rate of 0.5 ml/min in a Biorad NGC chromatography system. Fractions of 0.5 ml were collected. The column was calibrated with the following standards: Aldolase (158 kDa) Ferritin (449 kDa) and Thyroglobulin (669 kDa). The exclusion volume (Vo) was determined with dextran blue. Twenty-two fractions of 0.5 ml starting from the excluded volume were analyzed: 80 μl of each were mixed with 40 μl of 2-times concentrated Laemmli loading buffer, denatured at 100°C and 25 μl of the mixture were loaded onto 8% (for TRAPPC11-HA_3_ or Trs85-HA_3_ detection) or in 12% (for Trs23-HA_3_ detection) SDS-PAGE gels, transferred to a nitrocellulose membrane and analyzed by western blotting using anti-HA Roche rat monoclonal 3F10 (1/1000) and HRP-coupled goat anti-rat (Southern Biotech, 1/4000) as primary and secondary antibodies, respectively. HRP signal was detected with Clarity western ECL substrate (Biorad). Chemiluminescence was recorded with a Biorad Chemidoc Imaging System and the signal in anti-HA-reacting bands was quantified with Image Lab 5.2.1 (Biorad) and plotted with SigmaPlot using the ‘multiple Spline Curve’ option of the Graph menu.

#### S-agarose purification of TRAPPs under N-sufficient and N-starved conditions

Strains expressing Bet5-Stag or TRAPPC11-Stag were cultured for 14–15 h at 30°C on MFA medium before harvesting the mycelia by filtration and dividing the resulting biomass into two halves. One half was freeze-dried. The second was transferred to a minimal medium without nitrogen and incubated for 1 hour at 30°, filtered and freeze-dried. 1.75 grams of each freeze-dried sample was ground with ceramic beads in a Fast Prep. The powder was resuspended with 45 ml of extraction buffer [25mMTris-HCl, pH 7.5, 0.5% v/v NP40, 200 mM KCl, 4mM EDTA, 1mM DTT, 5 μM MG132, Complete ULTRA EDTA-free protease inhibitor cocktail (Roche)]. Then 5 ml of 0.6mm glass beads were added and the mixture was homogenized with a 15 sec full-power pulse of the FastPrep followed by incubation for 10 min at 4°C in a rotating wheel. This step was repeated two additional times before centrifuging the resulting homogenate at 15,000 x g for 30 min at 4°C and collecting approximately 35 ml of supernatant, which was spiked with 1% (w/v) BSA.

Then, 0.5 ml of S-protein agarose beads equilibrated with extraction buffer containing 1% BSA was added to each extract. The mixture was incubated for 2 hours at 4°C in a rotating wheel. Beads were collected by centrifugation at 4°C at 300 g in a tabletop centrifuge and washed four times for 10 min at 4°C with 10 ml of washing buffer (25mMTris-HCl, pH 7.5, 300 mM KCl, 4mM EDTA, 1mMDTT). For protein elution, the resin was transferred to a 2 ml Eppendorf tube and resuspended in washing buffer containing 4 mg/ml of S-peptide. The mixture was incubated at 37° for 15 minutes with gentle agitation, and the supernatant was collected by centrifugation. This step was repeated a second time. The eluate was precipitated with 10% TCA and the pellet was resuspended with 60 μl of Laemmli buffer. Half of the sample was loaded onto a 10% SDS-polyacrylamide gel that was run until proteins moved ∼ 1 cm into the separation gel. The portion of the gel containing the protein mixture (identified after staining with colloidal Coomassie) was excised and processed for MS/MS analysis.

#### GFP-trapping and MS/MS

Conidiospores from *wt* or *trs33*Δ strains were inoculated in flaks containing MFA, which were incubated overnight at 30°C. The mycelium was then filtered and transferred to nitrogen-free minimal medium for 1 hour at 30°C, filtered and freeze-dried. Cell extracts were prepared as described above, but using lysis buffer containing 25 mM Tris-HCl pH 7.5, 150 mM NaCl, 0.5 mM EDTA, 0.5% NP40, Complete ULTRA EDTA-free protease inhibitor cocktail (Roche) and 5 μM MG132. 4 mL of extract (approximately 100 mg of total protein) were immunoprecipitated with 25 μL of GFP-Trap magnetic agarose beads (Chromotek) for 2 hours at 4°C in a rotating Wheel. Beads were washed four times with the same buffer. For MS/MS analysis, the proteins immobilized on the resin were digested ‘on-beads’: The beads were resuspended in 25 μL of buffer I (25 mM Tris-HCl pH 7.5, 2 M urea, 100 ng trypsin (Pierce Trypsin Protease MS-Grade, Thermo Scientific), and 1 mM dithiothreitol, and incubated in a thermomixer at 37^0^C and 400 rpm for 30 minutes. The beads were pelleted on a magnetic stand and the supernatant was transferred to a fresh tube. Pelleted beads were resuspended in 50 μl of elution buffer II (50 mM Tris-HCl pH 7.5, 2 M urea, 5 mM iodoacetamide), and pelleted a second time to recover the supernatant, which was combined with the first one. This step was repeated a second time before combining the supernatants and incubating them in a thermomixer at 32°C and 400 rpm overnight protected from light. The digestion was terminated by adding 1 µl of trifluoroacetic acid and the sample dried in a speed vacuum centrifuge. The resulting peptides were purified on a Zip Tip with 0.6 μl of C18 resin (Merck), dried and resuspended in 10 µl of 0.1% formic acid in water. Four microliters of each sample were subjected to nano-LC ESI-MS/MS analysis using a Vanquish Neo nano system (Thermo Scientific) coupled to an Orbitrap Exploris 240 mass spectrometer (Thermo Fisher Scientific).

Samples were loaded onto an Acclaim PepMap 100 precolumn (Thermo Fisher Scientific), and eluted in a RSLC PepMap C18, 50-cm long, 75-µm inner diameter and 2-µm particle size. The mobile phase flow rate was 250 nL/min using 0.1% formic acid in water (solvent A) and 0.1% formic acid and 80% acetonitrile (solvent B). The gradient profile was set as follows: 4% solvent B for 2 min, 4–31% solvent B for 93 min, 31%-50% solvent B for 7 min, 50%-100% solvent B for 1 min and 100% solvent B for 15 min. For ionization, a liquid junction voltage of 1900 V and a capillary temperature of 275°C were used. The full scan method employed a m/z 350–1200 mass selection, an Orbitrap resolution of 60,000 (at m/z 200), a normalized target automatic gain control (AGC) value of 300 % and maximum injection time in automatic mode. After a survey scan, the 20 most intense precursor ions were selected for MS/MS fragmentation. Fragmentation was performed with a normalized collision energy of 50 % and MS/MS scans were acquired with a starting mass of m/z 120, AGC target was custom, resolution of 15000 (at m/z 200), intensity threshold of 1.0e4, isolation window of 1.0 m/z units and maximum injection time in automatic mode. Charge state screening was enabled to reject unassigned, singly charged, and equal or more than seven protonated ions. A dynamic exclusion time of 10 sec was used to discriminate against previously selected ions.

#### MS/MS data analysis

MS data were analyzed with Proteome Discoverer (version 3.0.0.757) (Thermo Scientific) using the Mascot search engine (version 2.6, Matrix Science). Searches were performed against Aspergillus nidulans FGSC A4 version_s10m02-r03_orf_trans_allMODI. (10,489 protein sequence entries). Precursor and fragment mass tolerance were set to 10 ppm and 0.02 Da, respectively. Trypsin/P was selected as the protease, with a maximum allowance of two missed cleavage sites, Cys carbamidomethylation as fixed modification, with N-terminal acetylation, Asn or Gln deamination, methionine oxidation and N-terminal pyrrolidonation of Glu and Gln as variable modifications. Identified peptides were filtered using Percolator algorithm with a q-value threshold of 0.01. For Fig 3 the abundance of each protein hit was given as the number of PSMs for each TRAPP protein divided by the number of Lys + Arg residues present in the protein sequence and multiplied by 100. For Fig 2J and K involving parallel purifications from wt and mutant cells these scores were plotted as % relative to the wt.

