## Supplemental Figures for "A TRAPPC2L/TRAPPC11/12/13 subcomplex directs TRAPPIII to autophagy"

### Supplemental Figure legends

**Figure S1.** Localization of GFP-tagged TRAPPC11 and TRAPPC12 to secretory compartments and to the PASs. The latter are indicated by arrowheads.

**Figure S2.** Four-way amino acid sequence alignment of human, *Drosophila*, *Arabidopsis* and *Aspergillus* TRAPPC11. The *foie gras* domain is shaded in purple. The two regions that Galindo et al determined as being involved in the interaction with TRAPPC2L are boxed in yellow. Similar amino acids were according to the Blosum 45 score table.

**Figure S3**: Top, four-way amino acid sequence alignment of *Aspergillus*, human *Drosophila* and *Arabidopsis* TRAPPC2L/Tca17. The region containing the critical Leu-Asp-Φ motif is boxed (Φ indicates a hydrophobic residue). Similar amino acids were according to the Blosum 45 score table. Bottom, AlphaFold prediction of the interaction between the *foie gras* domain of *Aspergillus* TRAPPC11 and Tca17. Left, overall view of the predicted complex. Right, details of predicted interactions between the relevant region of the TRAPPC11 *foie gras* domain with α-helix 1 in Tca17. Asp48 within the conserved Leu-Asp-Φ motif corresponds to Asp37 in both *Drosophila* and humans.

**Figure S4.** AlphaFold predictions of the complex formed by the C-terminal half of TRAPPC11 with TRAPPC12 and TRAPPC13. (A) Predictions for the indicated complexes shown in surface and cartoon representations. (B) Structural alignment of TRAPPC13 from the three species showing the loop that is located between two beta-sandwiches. Note that *Aspergillus* TRAPPC13 contains only two beta-sandwiches, whereas the human and *Drosophila* proteins contain three. (C) AlphaFold prediction of the loop of *Aspergillus* TRAPPC13 inserted through the central channel of the TRAPPC12 alpha barrel.

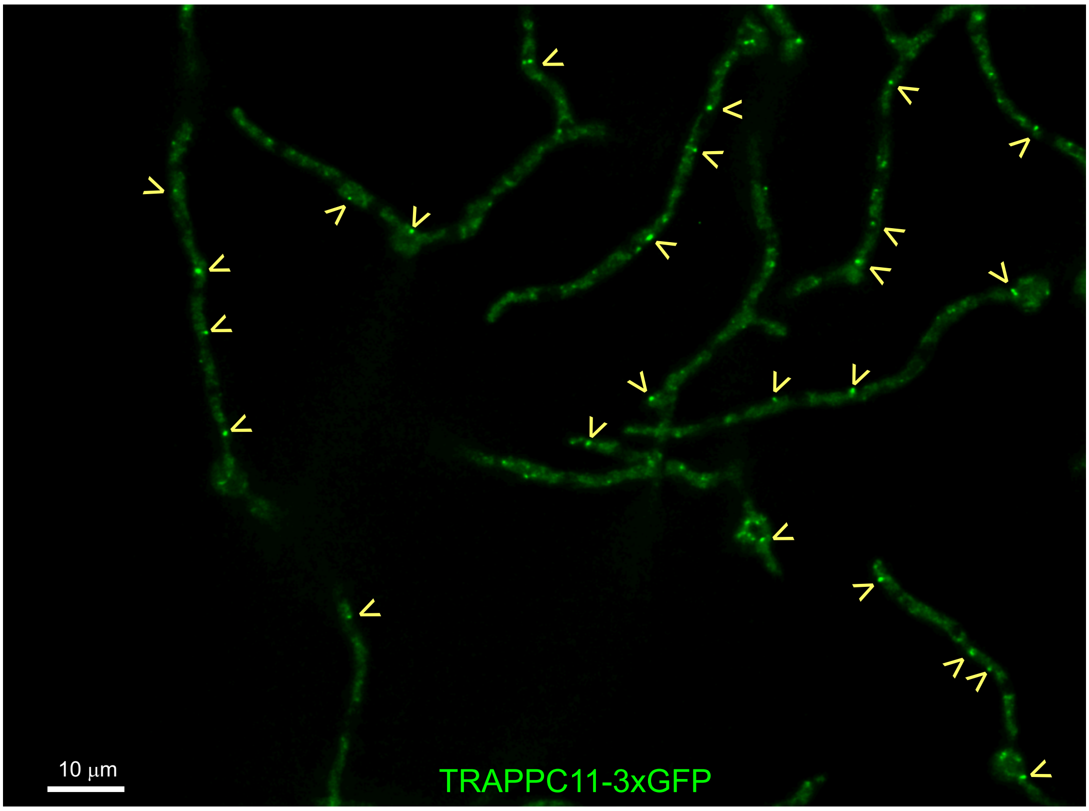


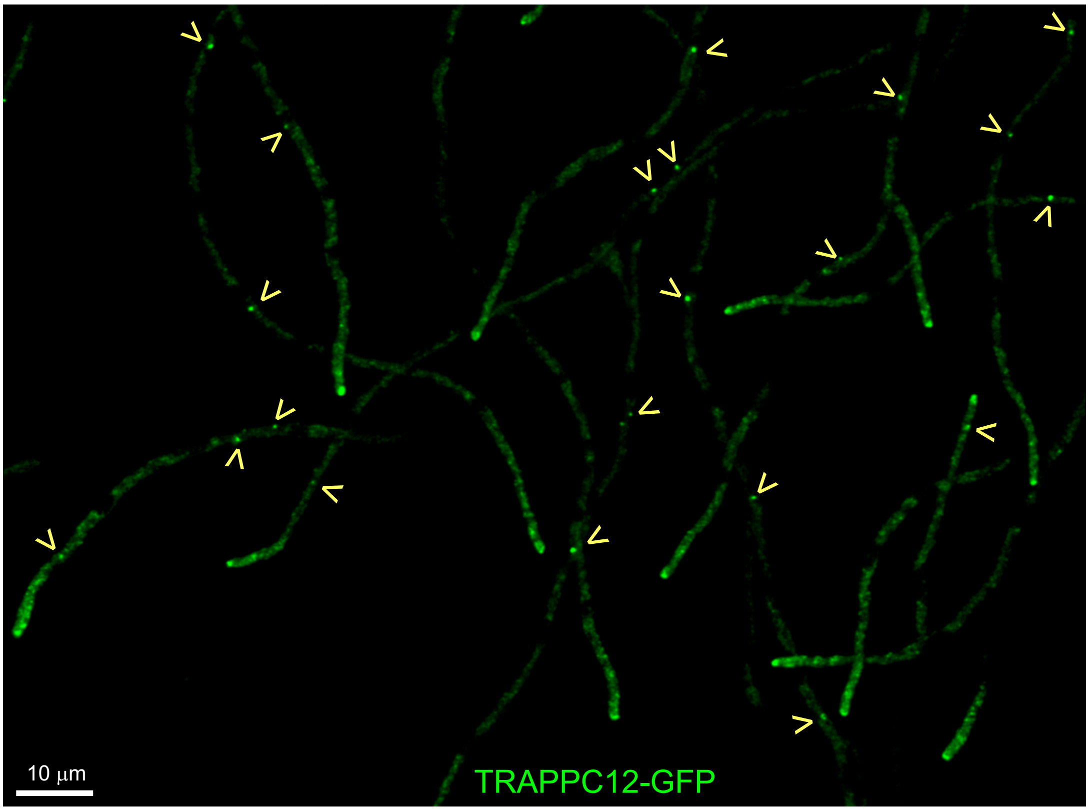


Figure S1

```
                *        20         *        40         *        60         *        80         *       100         *       120         *       140         *       160
HS : MSPTQWDFPVELCCRPMAFVTLTGLDVVYNAVH--------------------------RAVWDAFCAN--------------RRADRVPISFKVLPGDHEYP------------------------------KCRPKRTSYEWYIPKGILKTGWMNKHL :   90
DM : MTMDATALPSELLVTPQPLVGFCGLDTARVSVH--------------------------KAVWEAFSGSLQR-----------KAADRAAVQYKLLPPNYEFP------------------------------VAKPKRASYEWYHPKGILKRNWMLKHL :   93
AT : ME----EYPEELRTPPVSLVALFGYAELHASITKY---------------LHSQQPPINALAFPDFSQISLLL----AHDDQISRT-----------SSFRDP------------------------------LSVSDSASPIPSRCGGILKRDWLLKHR :   96
AN : MD----DYPRPYIANNLPFILLCGLEADGDRTEPEAEETNYPLLEENGIAIESDLPPLERPVAEEFRDVFFSEAGPRSQDNGIEGTSRAGYKVKSIGRNYRLPPRKANPPPASPPESPTVSRDHGLSMILHSPISPLTPA--SPTFPDGIMTPLWASKHQ :  154

                *       180         *       200         *       220         *       240         *       260         *       280         *       300         *       320
HS : NLVPALVVVFYELD---WDEPQWKEKQSECATRVEIVRQS--LQGRNTKVAVVLIQKKTPLPPGEDVIA-SERAAALCNACELSGKSLFVLPH--TD-HLVGYIIRLENAFYEHAQTYYYTEIRRVKSHKEFLN------------K--TTHQLLFVRHQ :  227
DM : HVLPSVVVLFQDME---WNDLQWTEKQVQCAAIVQALKNT--LQERNTRLCLVLLQRAAPLPPGEDLLA-AERAASLTNACGITSKMLFILPH--TE-HLTGYALRLESAFLDMAQSYYALMSKRIRNHRDQLT------------A---AHTSLKIRHQ :  229
AT : TKVPALVAAFFPSHHIFGDPTQWL----QVCSDLDSLKSV--IRPKNIKLVVVVVQS-S---PHEDIS--DDRLVALRKRAELDSKYVLFFNSSIVS-ELTLSLSRLASAFAELALSYYREEGRRIKSRIEKRS---------------SNSLDLNVRYC :  228
AN : DLVPAAVINFFPFC---LDSTMNSLRDNQLKIEINGLKKQWSSSGYRTRFIVVLLSEKA---IGDDLGDLDDRVLGIRRATNLDQKSLFILPPDVSSEEMRDFTRSLLALLHPSLMEYYRDLSKHARRKRNRSSIPPPTAPPTSGTSQTLSLQGWNTRYE :  308

                *       340         *       360         *       380         *       400         *       420         *       440         *       460         *       480
HS : FKIAFFSELKQDTQNALKNYRTAYNLVHELRAH------ETNILEIKTMAGFINYKICRLCFQHNTPLDAIAQFRKHIDLCKK---KIGSAELSFEHDAWMSKQFQAFGDLFDEAIKLGLTAIQTQ------------------------------NPG :  347
DM : FKLGFVAEMRQDFSTGQKHYFQAYANLDEIRIN------DGNCLEIKTLAGFLNYKICRLMFKLKTPRDAINQFIIHVEKHKS---RVGFKDLAFEHHAWLSTQHSVFAELFCEAIKNGLPALQTQ------------------------------HPG :  349
AT : FKVAVYAEFRRDWGEALKFYEDAYHSLHEMIGTSTRLPAIQRLVEIKIIAEQLHFKISTLLLHGGKLIEAVTWFHQHKTSYEK---VVGSTEFIFLHWDWMSRQFLVFAELLETSSATGQSLT-SSNSQGT-A--------------EISL-TEFEFYPA :  367
AN : FKMGIFAEFRQEIDAALKNYESAYETLFGQEVFENIAGWDPKFNEARLFSDALAIRIIRCLLSTGHTSSAVRSWVYHRRRTQDIVNRRGKGTRNYGWEAWEARWSSVMAQLIHQAKIPHLTPLETSQNQVHEHNSIFVPPEKTFLSVDEVLPWERLHHEG :  468

                *       500         *       520         *       540         *       560         *       580         *       600         *       620         *       640
HS : FYYQQAAYYAQERKQLAKTLC-NHEASVMYPN-PDPLETQTGVL-DFYGQRSWRQGILSFDLSDPEKEKVGILAIQLKERNVVHSEIIITLLSNAVAQFKKYKCPRMKSHLMVQMGEEYYYAKDYTKALKLLDY--VMCDYRSEGWWTILTSVLTTALKC :  502
DM : IYYHKAAEFVMKRRDAAMEAY-AAMQASSEAT-PTPIQNPLSLYTEFFGIRAVKT----GDLVA---EQQANMQLCDQERSYNHSAAIIALLSQAMAQFKIYKCLRFRKKLAIDMAEEYLKSGDHAKALTLYSL--MLPDYRQEKWTTIFTDVLLKTLRC :  498
AT : YYYQLAAHYLKDKKS-ALELLLSMSEIAQEIDSSSASITPSVYVGQFAQLLEKGEAITLHSITD---EEYTRYTI-SEAKRVQDSLQIIAWLKRSYESFTNLKAQRMAALCAFEVAREYFDLADPNNAKFFFDI--AANLYRQEGWVTILWEVLGYLREC :  520
AN : YWLYRSAKHTMHRRTLAGQIP-AEDR-------MPPGQSPASHIASKAYLYDT-------YLAP----ETHIEAARAQDGGVDHSTLILDTLKAALVEFSKRNQVRMTESLSLEIAEEYMRLGSWIEAYETLRPLWPALTWRRSGWWLIMAKFAWILREC :  609

                *       660         *       680         *       700         *       720         *       740         *       760         *       780         *       800
HS : SYLMAQLKDYITYSLELLGRASTLKDDQKSRIEKNLINVLMNESPDPEPDCDILAVKTAQKLWADRI-----------SL----AGSNIFTIGVQ---DFVPFVQCKAKFHAPSFHVDVPVQFDIYLKA---DCPHPIRFSKLCVSFNNQE--------- :  632
DM : ALLSGSVADYIACSVEALSLRHQSDQSERILILENLWQVFQGVPPMPKTQL----TPEAQALWTSAL-----------AN----VKSP-IQIDLD---KVNDVVEMCATFERVQLSNDDLLQLQLIVRV---LTDIPLRIRSFHVILADAGNP---QNSY :  629
AT : SRNLDALKDFVEFSLEMVALPVTSYENSGNLRNKNYG---PGGPATISGRES-----IHQEVFTLVCREAELLSSTEGSGFKLATDSP-LHLEIDLVSPLRPVLLASVAFHDQMIKPHALCSFTLSLLS---HLPLPVEIDHLEVQFNQST--------- :  659
AN : AARAQDSETVLRVDWELLNFAFPPRPD--------------------WNYD-----IHQSLAEFAS-----------KE----QKPS-IVIKAE---DVVSSVTATFIFQKAEGNVGEPLQCQLVLRSCAQKSSVPVRFAELRVAFEGCLRPIRLQSDQ :  724

                *       820         *       840         *       860         *       880         *       900         *       920         *       940         *       960
HS : -----YNQFCVIEE-------ASKANEVLENLTQ---------GKMCLVPGK---------------------------TRKLLFKFVA----------------------KTEDVG--------KKIEITSVDLALGNETGRCVVLNWQGGGGDAASS :  713
DM : KLEA-LKYFCFPTLTQLRGQKQP-DDEQLENPSQEPKNFE---KNMRLEPGS---------------------------YYQLFCSTEA----------------------QQFHEN--------TQLRIVRLEAHMGTDQVAALL---------TCSS :  717
AT : -----CN-FV-I-RNSQRPLWASA-SNTV-KSGS---QVE-NAPLLVLVPNN---------------------------WLRLTYAI------------------------KSEQS---------GKLECLSVLAKLGPLFTICSR-------AESPAA :  737
AN : NIDADTSTTCLISSPSLRDPSATS-DSSVHQSTTSALNALTGIADLTLGPAQTKVYNLACIPRESGESRVASIALLVSQEDFDLTYAITDLAQQEAFWWQQTTKGPARRRVGKGRDVDRCKIQPKPPKIRLTIPNLKSTYYTNERIVLNLVIDNGEDEA- :  882

                *       980         *      1000         *      1020         *      1040         *      1060         *      1080         *      1100         *      1120
HS : QEALQAARSFKRRPKLPDNEVHWDSIIIQASTMIISRVPNIS---------VHLLHEPPALTNEMYCLVVTVQSH-EKTQIRDVK-----------------------------------------------LTAGLKPGQDAN---LTQ-KTHVTLHGTE- :  813
DM : NYSRQLFRHHTRSRDLDDN-VTINPICYIAPTFHLDTQTNLGHGHDNGLTDDKEMVATKMLVNEYFPVVTTVSNP-YNVYLQNVGVHISIPVGLRNSVFLTTDISPGRQKLHSQIQIDVGELSAHGSNTATFYIFSLTEAEIKLAQRL-SYTLDVDRGPG :  874
AT : MEDLPVWKHENRVESLPTKDP-VLAVFGQKATQVDEPEPQVD---------VSLGASGPALVGEDFAMPIVVTSKGHAVYSG-------------------------------ELKI-------NLVDVGGGGLFSPREAEP---FSLESHHVEICGIDG :  846
AN : ADVVAEARLFG-----------------------------------------------------------SPDS-------------------------------AVNI-------SWLDQ-EGSVQASTESRP------------------ :  918

                *      1140         *      1160         *      1180         *      1200         *      1220         *      1240         *      1260         *      1280
HS : -----------LCDESY---PAL---------LTDIPVGDLHPGEQLEKMLYVRCGTVGSRMFLVYVSYLINTT-VEEKEIVCKCHKDETVTIETVFPFDVAVKFVSTKFEHLERVYADIPFLLMT------------------------------------- :  915
DM : GGGSNARVSTAPNSSESTPDHEVKSVIPSVAAISPVQIEYLD------------ETR-------------------------LRKSREDTLTVRCYGDFKFNARFYTMDRKPLNQVYRGENFLLRANTEVVAVDDVEILDSFFICDHNLVQSNYSFKQKK :  997
AT : A---------EGNNESESETGSIKKI---QQSFGLVSVPYLKEGESWSCKLEIKWHRPKPVMLFVSLGYLPHGS-EANTQ---KVHIHKSLQIEGKMPLLISNRFM--------LPYRRDHLLL-------NRIKPAPDSE-------DVSSLP------ :  962
AN : ------------NTPAEEEPHSLKRSVGVIESSSQREIPIVIS------------GTESSFEYELEVSAAYNLFSDIQT----PIIATTRLTLPIIRPFEANYEF------------------L-------PRLNPLPWPDFFTIDDDLTETQP------ : 1019

                *      1300         *      1320         *      1340         *      1360         *      1380         *      1400         *      1420         *      1440
HS : ----------------------------------------------------------------------------DLLSASPWALTIVSSELQLAPSMTTVDQLESQVDNVILQTG----------------ES-----ASE----------------- :  961
DM : YTNKYSAGEQLESVIVLRTNATLRDWATA--RDLDHRGKLEGKAVASKFIRPVPKPSSEEPLAPTPPAGISAYMSKSLVTKIPTT--------------MTIINSNSAVSNALQVANAGVMSNSTQSDDVNVAEE-----AINTHEGIKTTRLIYNKALE : 1136
AT : -------------------------LNEKSVLVVS------------------------------------AKNCSEIA------LKLVS---------MSIEFDDEQGETSCLIQQGGGCGDSPSSANLAPGEE-----FKKVFTVIPTTR-------- : 1033
AN : -------------------------NTEP--RGLQQRWCLDTK----------------------------VVSFAREP------LVIEN---------MSIAVLSLSG--GAISQVGHEVLVSPETREIHPEDLRHSNFVLDIQKIILGDR-------- : 1099

                *      1460         *      1480         *      1500         *      1520         *      1540         *      1560         *      1580         *      1600
HS : ---------CFCLQC-PSLGNIEGGVATGHYIISWKRTSAMENIP-----IITTVITLPHVIVENI-PLHVNADLPSFGRVRESLPVKYHLQNKTDLVQD--VEISVEPSDAFMFSGLKQIRLRILPGTEQEMLYNFYPLMAGYQQLPSLNINLLRFPN- : 1102
DM : AVQGTGHCRGFIKKIYTLEGNPSAVPIFGVFCIRWRRANCKEE--------NESKFVIRGLDIAEP-PLNIYCTIEEKMFVKMPMAFKVVLKNPTTHVLHLIANLSISKTDNFICSGHKQLDISIMAYEEKELVYNLYPLQVGWQELPVLSLEYNTKADP : 1287
AT : ----------------------TPKLGLGSIHLKWRREGGNITEA-----YVSTKHKLPEVNVEAS-PLVMSLDSPPYAILGEPFTYAVRICNQTQLLQE--AKFGLADAQSFVLSGSHSNTVSVLPKSEHVLSYKLVPLTCGEQQLPKITLTSAR---- : 1159
AN : ----------------------RPTSLNLALEIQWRRSGEETDDSGRSDRLTTTKLAIPRFVIPAGEPRVLASAISSR-KFPGLIHVEYTLENPSLHFLT--FSLTMEASEYFAFSGPKTMVVQLAPVSRQTVRYNLLASKRGLWIQPQLLVV------- : 1227

                *      1620         *      1640
HS : --------FTNQLLRRFIPTSIFVKPQGRLMDDTS-----IAA-A : 1133
DM : QKKDSQNALLDELVQRALPKRVFVLPPLKQQN------------K : 1320
```

Figure S2

```
             *        20         *        40         *        60         *        80
AN : MPGPKIACIGVIGKASDRLKDNPLHISLFPPYLGSAIDFSFLLNSSLDIFEIRQK------QTSVDQDLGLLQAVDERLASYGWITTTG :  83
HS : MA----VCIAVIAKE-----NYPLYIRS-T-PTENELKFHYMVHTSLDVVDEKISAMGKALVDQRELYLGLLYPTE-DYKVYGYVTNSK :  77
DM : MA----FCIAVIGKD-----NAPLYLTT-S-DMEQELELQYHVNAALDVVEEKCL-IGKGAPESKELYLGLLYSTE-NHKIYGFVTNTR :  76
AT : MI----VCVAVVGHQ-----NNPLYIQSFT-DADDALKLHHIVHCSLDVIEERVNNPKKSGTTLNEAFLGLLYPTV-NYKVYGYLTNTK :  78

       *        100         *       120         *       140         *       160         *       180
AN : LKFLIIVDLIGDQTSSVVNEVRGSVRAGVREADLKPAFRALQAAYIRLLQNPFYSPDDHAIINGAVQGTRPSQIT-DQRFIAEVNRIGN : 171
HS : VKFVMVVD---------------SSNTALRDNEIRSMFRKLHNSYTDVMCNPFYNPGDR--------------IQSSRAFDNMVTSMMI : 137
DM : VKFIVVID---------------SSNVALRENEVRAIFRNLHLLYTDAICNPFYIPGES--------------LT-SKKFDRAVQKLMS : 135
AT : VKFILVT----------------TDLDVRDTDVRSFFRKFHAAYVDAVSNPFHVPGKK--------------IT-SRTFAQTVSNIVG : 135

AN : SWGAVTSNV : 180
HS : QV--C---- : 140
DM : GT--A---- : 138
AT : SYGLN---- : 140
```

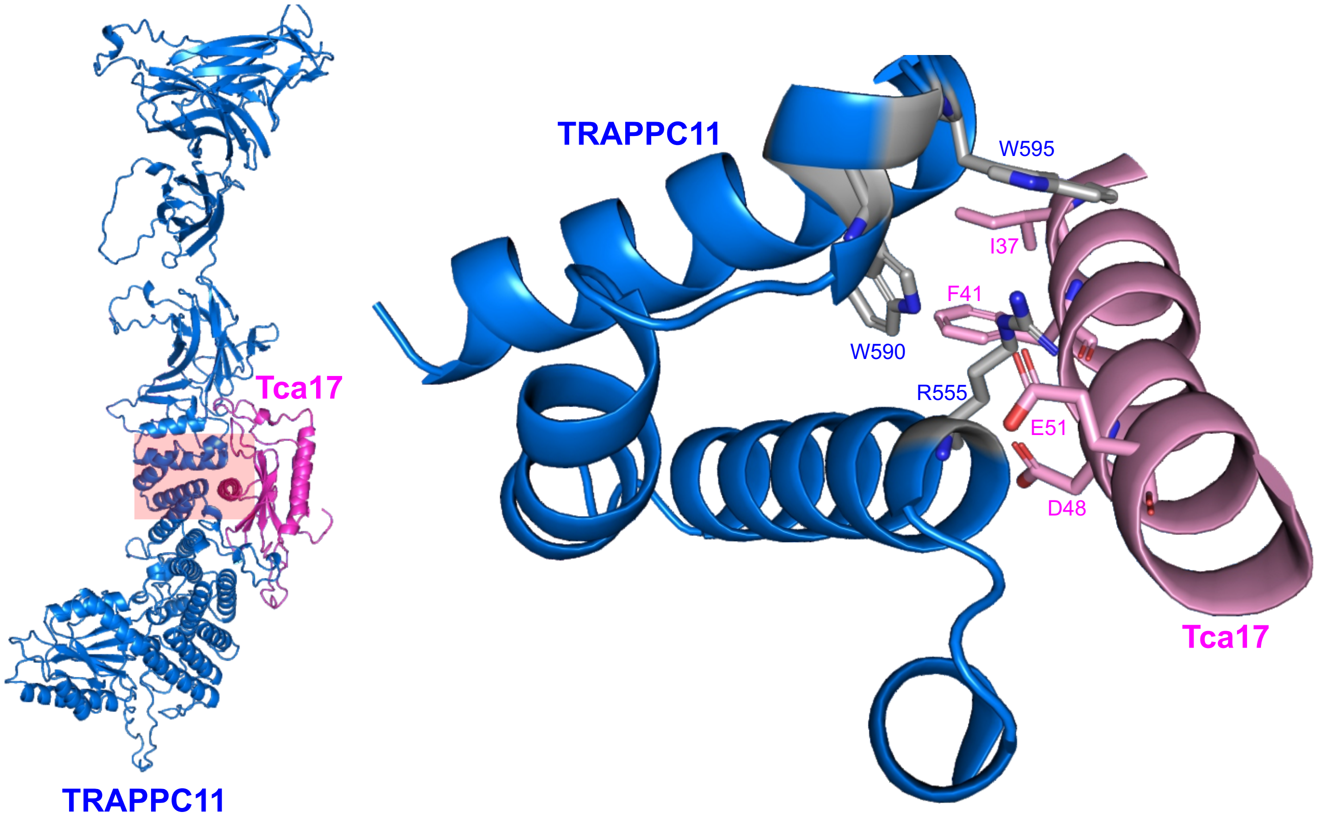


Figure S3

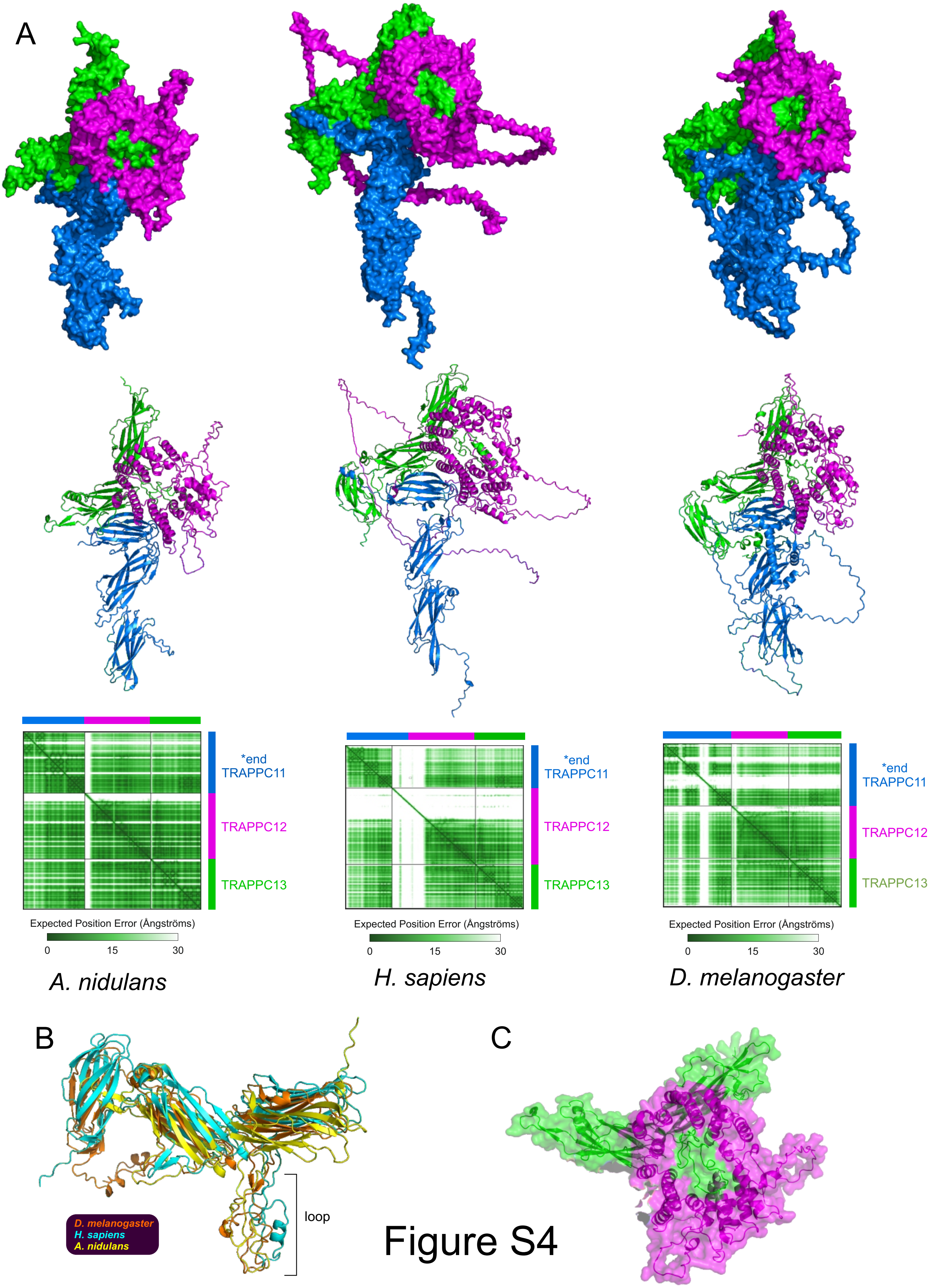
A
*end
TRAPPC11
TRAPPC12
TRAPPC13
Expected Position Error (Ångströms)
0
15
30
A. nidulans
*end
TRAPPC11
TRAPPC12
TRAPPC13
Expected Position Error (Ångströms)
0
15
30
H. sapiens
*end
TRAPPC11
TRAPPC12
TRAPPC13
Expected Position Error (Ångströms)
0
15
30
D. melanogaster
B
D. melanogaster
H. sapiens
A. nidulans
loop
C

Figure S4
